# Genomic analysis of the probiotic candidate *Bifidobacterium bifidum* strain 900791 in the context of the *B. bifidum* pangenome

**DOI:** 10.1101/2025.11.10.687612

**Authors:** Juan P. Cárdenas, Boris Vidal-Veuthey, Kevin Meza, Daniel E. Almonacid, Anastasia Gutkevich

## Abstract

**Background:** *Bifidobacterium bifidum* is a key commensal bacterium in the human gut microbiota with recognized probiotic properties and health benefits. An underinvestigated strain called 900791, isolated from meconium of a Siberian infant and with a validated role in lactose tolerance, requires more understanding of the genomic basis of its potential probiotic functionality.

**Methods:** We performed whole genome sequencing of strain 900791 using hybrid sequencing (Illumina NextSeq and Nanopore). The genome was functionally annotated and compared against 228 *B. bifidum* genomes to elucidate probiotic determinants and evolutionary relationships. Antibiotic resistance profiling was conducted using three independent approaches, carbohydrate-active enzymes (CAZymes) were characterized using dbCAN analysis; additionally different markers for desired phenotypes in probiotics (such as adherence to epithelial cells, resistance to low pH, bile salts and oxidative stress, and the ability to produce bacteriocins) were also investigated.

**Results:** The complete genome comprises 2.28 Mb with 62.43% G+C content, encoding 1,852 ORFs, 53 tRNAs, and three complete 16S rRNA genes. Phylogenomic analysis revealed 900791 belongs to a distinct clonal subgroup of nine closely related strains sharing almost identical allelic similarity (observed via cgMLST). The pangenome analysis of 229 *B. bifidum* genomes showed an open structure with 4,679 orthogroups, including 1,212 core families.The strain 900791 harbors extensive CAZymes from families GH2, GH20, GH33, and GH84 associated with mucin and human milk oligosaccharide degradation, plus GH2 and GH42 families for lactose metabolism. Safety assessment revealed only intrinsic resistance to mupirocin and rifampicin, with no acquired resistance markers or virulence factors. The strain possesses multiple stress tolerance systems including acid resistance (F₀F₁-ATPase), bile salt resistance (MFS transporters, BSH hydrolase), and oxidative stress response mechanisms. Predicted probiotic features include adhesion proteins (FimA/FimB, FimM), bacteriocin production capabilities, and comprehensive stress response systems.

**Conclusion:** *B. bifidum* 900791 harbors markers for significant probiotic potential with genetic features optimized for gut colonization, host interaction, and antimicrobial competition. This genomic characterization provides a framework for understanding *B. bifidum* diversity and supports *B. bifidum* strain 900791-specific probiotic applications, particularly given its demonstrated efficacy in improving lactose tolerance in human subjects.

## 1. Introduction

*Bifidobacterium bifidum* is a key member of the human gastrointestinal tract microbiota, recognized for its primary roles in maintaining gut homeostasis, modulating immune responses, and shaping the microbial ecosystem, especially during human early life (Turroni et al., 2019). Several strains of this bacterium have been extensively used in the elaboration of different foods such as cultured milk and other dairy products (Yaeshima et al., 1996). *B. bifidum* strains have been also utilized in therapeutic preparations for the treatment of digestive disturbances in infants (Bellomo et al., 2024; Totsu et al., 2014). *B. bifidum* exhibits the metabolic capacity to degrade host-derived glycans, including human milk oligosaccharides (HMOs) during its colonization in infants, as well as mucin glycans in the adult gut (Xiao et al., 2024). This feature enables this organism to colonize and persist throughout different human life stages. *B. bifidum* supports the integrity of the intestinal epithelial barrier by stimulating mucin production and enhancing the protective mucus layer that limits pathogen invasion and maintains intestinal integrity (Martín et al., 2020; Xiao et al., 2024). Additionally, the glycan-degrading ability of *B. bifidum* into simpler sugars not only sustains its own growth but also facilitates cross-feeding interactions with other gut microbes (Xiao et al., 2024). This syntrophy promotes an eubiotic microbial community and increases the production of short-chain fatty acids (SCFAs), such as acetate and butyrate, which are considered beneficial substrates for gut health and epithelial function (Morrison and Preston, 2016; Parada Venegas et al., 2019). Interestingly, *B. bifidum* strains can also stimulate mucin synthesis in an indirect fashion, strengthening the mucus barrier and reducing intestinal permeability, which is crucial for protecting the host from pathogenic challenges (Engevik et al., 2019). Different studies suggested that *B. bifidum*, often in combination with other bacteria, can also influence the gut microbial community by fostering the growth of beneficial microbes while suppressing inflammation and potential pathogens (Shin et al., 2024). Its presence has also been linked to improved bowel function and relief from constipation, possibly through modulation of gut neurotransmitters (Fuyuki et al., 2021; Makizaki et al., 2021). In addition to its metabolic activity, *B. bifidum* exerts significant immunomodulatory effects: some studies suggest that this organism interacts with intestinal epithelial and immune cells, such as dendritic cells, to promote anti-inflammatory responses and support the maturation of the immune system, particularly during the early years (Lim and Shin, 2020). These interactions help maintain immune tolerance and prevent excessive gut inflammation, contributing to overall host health. The main sources of *B. bifidum* isolation reflect its ecological niche and metabolic specialization. It is predominantly isolated from the feces of breast-fed infants (Shah, 2011) or even breast milk (Martín et al., 2009). Human breast milk itself serves as a reservoir for *B. bifidum*, facilitating vertical transmission from mother to infant and establishing early colonization of the infant gut; therefore, their affinity for Human Milk Oligosaccharides (HMOs) can also be targeted for isolation (Kitaoka, 2012).

The rise of genomics, and more recently, the emergence of Next-Generation Sequencing (NGS) technologies, made a tremendous impact on microbiology, reflected in the generation and availability of massive amounts of genomic data, facilitating a variety of DNA sequencing approaches (Satam et al., 2023). Nowadays, whole genome sequencing (WGS), very useful for the generation of a first landscape of main metabolic and functional features for a bacterial strain, has been facilitated with hybrid sequencing techniques, combining the virtues and properties of short-(e.g., Illumina. DNBSEQ) and long-read (e.g., Nanopore, Pacbio) sequencing (Neal-McKinney et al., 2021). *B. bifidum* is just one of the several *Bifidobacterium* species, sequenced from a variety of different hosts, sources or geographical contexts (Alessandri et al., 2021). Recent genomic analyses of *Bifidobacterium bifidum* have elucidated its evolutionary strategies for gut colonization, emphasizing pan-genome plasticity and niche-specific adaptations. The first report for a genomic analysis for *B. bifidum* (the PRL2010 strain) showed the presence of genes encoding potential enzymes for mucin degradation such as extracellular sialidases (GH33), fucosidases (GH29), and lacto-N-biosidases (GH20) to degrade mucin oligosaccharides, enabling it to exploit mucin as a primary carbon source, confirming this feature by transcriptomic profiling (Turroni et al., 2010). Age-related genomic and phenotypic variation in *B. bifidum* was also explored: the comparison of six strains isolated from subjects in different age ranges suggested enhanced genomic capacity for utilizing 6’-sialyllactose, a key component of human milk oligosaccharides (HMOs), in strains from children and young people (0 - 17 years old) in comparison with adults (18 - 65 years old) (Wei et al., 2023). Another comparative genomics study (Abdelhamid and El-Dougdoug, 2021) analyzed 15 *B. bifidum* genomes from diverse human niches such as gut, vagina, and breast milk; this study showed the presence of differentially conserved carbohydrate active enzymes (CAZymes) reflecting different adaptation across different niches. A more recent study (Lu et al., 2021) compared 140 strains isolated from Chinese adults and infants, revealing an open pan-genome of 8,399 genes and a conserved core genome (638 genes). Phylogenetic clustering by geographical origin and isolation source (infant vs. adult feces) highlighted the influence of host ecology on genetic divergence. Notably, the study identified novel type III-A CRISPR-Cas systems in Chinese strains, suggesting localized adaptations to phage predation. Genetic diversity was pronounced in carbohydrate metabolism pathways, including glycoside hydrolases (GHs) targeting host-derived mucins, and antimicrobial competition factors like bacteriocin operons. Thus, the availability of *B. bifidum* genomic sequences may help to understand more of their abilities as members of the gut microbiota or as probiotic agents.

In this study we present the genome analysis of a strain of *B. bifidum* utilized in food preparations: the 900791 strain was isolated from the meconium of an infant from Siberia (Гуткевич et al., 2003). This strain has been tested successfully in the improvement of lactose tolerance in a human population cohort (Aguilera et al., 2021). We use genomic analysis and compare it with a current set of 228 taxonomically valid strains of *B. bifidum* to identify its genomic singularities, and understand its probiotic potential.

## 2. Methodology

### 2.1. DNA Isolation, Library Preparation, Hybrid Sequencing and Basic Annotation

The genomic DNA of this strain was extracted using the *FastDNA SPIN Kit for Soil* (MP Biomedicals) in accordance with the manufacturer’s instructions. The DNA was quantified using gel electrophoresis with 1% of agarose gel and fluorescence-based Qubit dsDNA DNA quantification System (ThermoFisher, USA). Hybrid sequencing of the isolate was carried out at EzBiome (Gaithersburg, MD, USA). Briefly, the genomic DNA was sequenced on an Illumina NextSeq2000 (2 × 150 bp) and an R10.4.1 flow cell of a Nanopore PromethION (Eugene, USA). The Illumina library was prepared using the NEBNext® Ultra™ II FS DNA library kit for Illumina, while the Nanopore library was prepared using v14 library prep chemistry without fragmentation or size selection. The resulting sequencing reads were filtered using Filtlong v0.2.1 (parameters: “--min_length 1000 --keep_percent 95”, see https://github.com/rrwick/Filtlong) by removing the 5% worst fastq reads. Nanopore reads were then assembled with Flye v2.9.2 (Kolmogorov et al., 2019). Illumina reads were aligned to the draft assemblies using BWA with the ‘-a’ flag (Li, 2013). Alignment files and draft assemblies were used to produce polished assemblies, using Polypolish v0.5.0 (Wick and Holt, 2022). Polished assemblies were checked for contamination using CheckM2 (Chklovski et al., 2023), annotated with Prokka v1.14.6 (Seemann, 2014) and circular genome maps were produced with GenoVi v0.4.3 using default parameters (Cumsille et al., 2023).

### 2.2. Antibiotic Resistance Gene Profiling

Antibiotic resistance gene profiles were evaluated using three different approaches. The first consisted in the use of a pre-built *bowtie2* (Langmead and Salzberg, 2012) database composed of NCBI’s *National Database of Antibiotic-Resistant Organisms* (NDARO, www.ncbi.nlm.nih.gov/pathogens/antimicrobial-resistance/) reference genes. Each read of the metagenome sample was mapped against these genes using *bowtie2* with the “--very-sensitive” option, and the output was then converted and sorted by *samtools* (Li et al., 2009). Finally, for each gene found, depth and coverage were calculated by using samtool’s *mpileup* script. The second approach consisted in analyzing the genome assembly by using ResFinder-4.7.2 (Florensa et al., 2022) using a cutoff of 90% identity and 60% minimum coverage. The third approach utilized the predicted ORFeome, which was analyzed by the *Resistance Gene Identifier* tool from the CARD database (*RGI-main* 6.0.3, CARD 4.0.0), considering perfect and strict hits only (Alcock et al., 2023).

### 2.3. Selection of *B. bifidum* genomes and generation of the phylogenomic tree

The following method differs slightly from those presented in a previous study (González et al., 2023). All genomes available in NCBI Genomes belonging to any taxonomic group associated to *Bifidobacterium* (NCBI:taxid 1678) were downloaded from the NCBI Genbank FTP site in early March 2025, and their metadata was extracted from their respective NCBI Project / Biosample accession (unless more curation were required). The taxonomic identity was confirmed by using the program ‘classify_wf’ of the GTDB-Tk program, version 2.4.0 (Chaumeil et al., 2022), using the database release 220 as the reference, selecting all genomes classified into the *B. bifidum* species (“s Bifidobacterium bifidum”). Genome completeness and contamination were calculated using the program ‘predict’ from CheckM2 (see above). Only genomes with the aforementioned taxonomic assignment, plus a completeness higher than 90%, contamination lower than 5% and an assembly N50 value equal or higher than 200 Kbp (see Results) were selected. The ORFeome of the resulting genome set was predicted using Prodigal, version 2.6.3 (Hyatt et al., 2010) (relevant parameters: *-q -c -m*). Using this ORFeome dataset (in addition to the 900791 strain), the orthogroup catalog was calculated by Orthofinder version 2.5.5 (Emms and Kelly, 2019) using “-S diamond_ultra_sens -y -og” as relevant parameters. To make the phylogenomic tree, a multiple sequence alignment was constructed from a set of 769 concatenated, single-copy conserved orthogroups by using MAFFT version 7.490 (parameters: --maxiterate 1000 --localpair) (Katoh et al., 2002); the alignment was used by *iqtree* version 2.1.4 (parameters: -m TEST --alrt 1000) to generate a maximum likelihood-based tree with an aLRT with 1000 replicates as the branch support test (Minh et al., 2020). The RefSeq ORFeome from *Bifidobacterium longum* subsp. *longum* JCM 1217 (GCF_000196555.1) was utilized as the outgroup. The phylogenomic tree was generated using the *ggtree* R package (Xu et al., 2022).

### 2.4. Pangenome Analysis

This following method also is a modified version from those presented in a previous study (González et al., 2023). The orthogroup matrix, obtained from Orthofinder results (including unassigned orthogroups), was utilized for different pangenome metrics. Pangenome curves were created using the *panplots* function in R (created by *SioStef*, https://github.com/SioStef/panplots/), using 1000 permutations. The □ value from the Power law deduced from the pangenome curve (Tettelin et al., 2008) was calculated using the function *curve_fit* from *scipy* python package, using the equation *“y = ax*^γ^*”,* applied on the *panplots* output. The shell, cloud, “soft-core” and core components of the pangenome were calculated by Python scripts with the *pandas* package, considering the following criteria: *core* gene families as the orthogroups present in 100% of the strains, soft-core groups as present in between 99.999 and 95% of the strains, shell as groups present between 94.99% and 15% of strains, and cloud as the gene families present in between 14% and 0.87% (the equivalent to two strains). Unique groups can be deduced from the set of “species-specific orthogroups’’, and the “unassigned genes,” both reported by Orthofinder. Heatmaps were created by using the *pheatmap* function from R. Figures were elaborated with *ggplot2* and the *ggarrangment* packages. In order to create core-genome multilocus sequence typing (cgMLST), the nucleotide sequences for the single-copy gene families were retrieved and an allele database was created by using pyMLST (Biguenet et al., 2023). The dataset of 228 genomes plus the 900791 strain were cataloged using this dataset.

### 2.5. Additional functional annotation processes

The predicted ORFeome from both the 900791 strain genome and the selected available *B. bifidum* genomes were analyzed by EggNOG-mapper (parameters: “--tax_scope_mode narrowest --tax_scope prokaryota_broad --go_evidence experimental”, see (Cantalapiedra et al., 2021)). Metabolic pathway prediction was conducted by the combination of the KAAS (Moriya et al., 2007), BlastKOALA (Kanehisa et al., 2016), and KofamKOALA (Aramaki et al., 2020) tools. KAAS was utilized in best-bidirectional-hit (BBH) mode, using *Prokaryote* set as reference; BlastKOALA was used also with the *Prokaryote* reference set; KofamKOALA was configured using 10^-6^ as the threshold E-value. Results were mapped by using *KEGG mapper*. Carbohydrate active enzymes (CAZymes) (Flint et al., 2012) were searched using the HMM database (v. 11) from dbCAN, the search tool based on CAZy (Zheng et al., 2023), using an e-value < 1e-10. Bacteriocins were predicted by using the BAGEL4 server (van Heel et al., 2018) using the assembly data. Subcellular localization predictions using the 900791 strain ORFeome were performed by PSORTb 3.0 for Gram-positive bacteria (Yu et al., 2010). When necessary, Diamond v. 2.1.8 (Buchfink et al., 2015) was used for BLASTP searches of the predicted ORFeome of the 900791 strain and other genomes against the Swissprot database (Release 2025_03).

## 3. Results

### 3.1. Overall properties of the genome of *Bifidobacterium bifidum* strain 900791

The hybrid sequencing of the genome of the *B. bifidum* 900791 strain resulted in a complete genome, consisting of a single circular chromosome of 2,280,092 base pairs (average depth: 22x) with an average G+C content of 62.43% (Supplementary Figure 1). The taxonomic status of this genome could be confirmed by two approaches. In the first approach, the predicted 16S rRNA gene from this genome exhibited 100% identity with the 16S rRNA gene from *B. bifidum* strains KCTC3202 and NBRC100015 (accessions NR_117505.1 and NR_113873.1 respectively). Using GTDBtk as a second approach, the identity of this genome as *B. bifidum* was confirmed by ANI against the genome of *B. bifidum* ATCC 29521 (GCF_001025135.1, ANI = 98.86%). Prokka annotation of this genome revealed 1,852 protein-coding genes (ORFs), 53 tRNA genes, and 6 rRNA genes. The protein-coding sequences account for approximately 85% of the total genome. This genome contained no plasmids. According to CheckM2, its contamination level is 0.12% (Specific model).

From the ORFeome of this strain, some general features can be predicted. Using PSORTb as the predictor of subcellular localization of the proteins, the analysis of the ORFeome of *B. bifidum* 900791 (Supplementary Figure 2-A) detected a set of 28.2% proteins outside the cytoplasm/cytosol, predicted to be located in the plasmatic membrane (n = 476, % 25.7), or the cell wall (n = 18, % 0.97); a total of 28 ORFs (% 1.51) were predicted to be secreted into extracellular medium. This may suggest that the ORF patrimony of this species could include potential secreted enzymes and effectors that deserve further attention. The functional annotation results from the EGGNOG mapper database (excluding 257 unclassified ORFs) showed that the proportion of sequences associated with *metabolism* (categories C, G, E, F, H, I, P, Q) represents 36.78% of the total assignments for this ORFeome (Supplementary Figure 2-B); functions associated with *information storage and processing* (categories J, A, K, L) represented 26.43% of all the CDSs, whereas functions associated with *cellular processes and signaling* (categories D, Y, V, T, M, N, Z, W, U, O, X) represented 19.3% of CDSs. Sequences from the metacategory *poorly characterized* (categories R and S) represented 17.49% combined. Analyzing by categories (Supplementary Figure 2-C), the most represented in the 900791 strain were associated with category S (“ *Function unknown*”, with 17.49% of assignments), followed by genes involved in *Amino acid transport and metabolism* (category E, with 9.18% assignments), *Replication, recombination and repair* (category L, 9.01%), *Translation, ribosomal structure and biogenesis* (J, 8.77%), *Transcription* (K, 8.6%) and *Carbohydrate transport and metabolism* (category G, 8.42% of assignments).

Since the absence of antimicrobial resistance is a very important trait in potential probiotics (Binda et al., 2020), the prediction of this feature was conducted by three independent strategies, one based on mapping against NDARO, other comparing against Resfinder and a third one based on RGI-CARD. According to NDARO and Resfinder, no antimicrobial resistance marker was predicted. However, predictions from RGI-CARD showed two classes of antimicrobial resistance: an intrinsic resistance to mupirocin, given by *ileS* alleles (ARO:3003730, 99.46% identity, 100.09% coverage) and an intrinsic resistance to rifampicin, given by *rpoB* alleles (ARO:3004480, 91.22% identity, 100.08% coverage). Those two resistance markers were driven by mutations previously reported to be part of the bifidobacterial genetic patrimony (Serafini et al., 2011; Lokesh et al., 2018). In contrast, specialized enzymes or acquired markers were not found in *B. bifidum* 900791. These results may suggest that this strain can be suitable for its use as a probiotic.

### 3.2. The 900791 strain gene content in the context of the *B. bifidum* pangenome

In order to obtain a better picture of the functionality of the 900791 strain, we also compared the predicted gene content of this strain with a set of 228 *B. bifidum* genomes, to obtain the pangenome of this species. According to metadata search (Supplementary Table 1), most assemblies were obtained from reported human samples (214 strains, 93% of the total). More than 79% of the selected assemblies were obtained from isolated strains. Only 30 metagenome-assembled genomes (MAGs) were found. Also, most strains were reported to be obtained from stool (60% of the total), several others without clear metadata (e.g., mentioning that are from faecal samples but not establishing whether they are from adults or children), and only few from other sources such as bladder, fermented milk, rumen or vaginal samples. We could obtain a phylogenomic tree from a set of 769 shared single-copy orthologs, using a strain of *Bifidobacterium longum* as the outgroup (Figure 1). Intraspecies tree clustering shows the presence of different clusters, suggesting the existence of different subtypes in the *B. bifidum* intraspecific diversity. Interestingly, the 900791 strain was strongly related of other eight genomes, belonging to the strains 791 (GCA_001595435, GCA_022014355), ICIS-310 (GCA_002114145), ICIS-643 (GCA_003790385), ICIS-202 (GCA_004799295), VKPM Ac-1784 (GCA_016070035), ICIS-629 (GCA_020710245), and ICIS-176 (GCA_020861855), all of them isolated from human stool samples (Supplementary Table 1). Moreover, the results of the cgMLST suggested that 900791 and the other eight genomes contained practically the same alleles (>99%) with each other, in comparison to the rest of the genomes. This phylogenetic pattern and allele content suggest that strain 900791 could be part of a clonal sublineage of *B. bifidum*, with potentially distinct genomic features.

**Figure 1.**
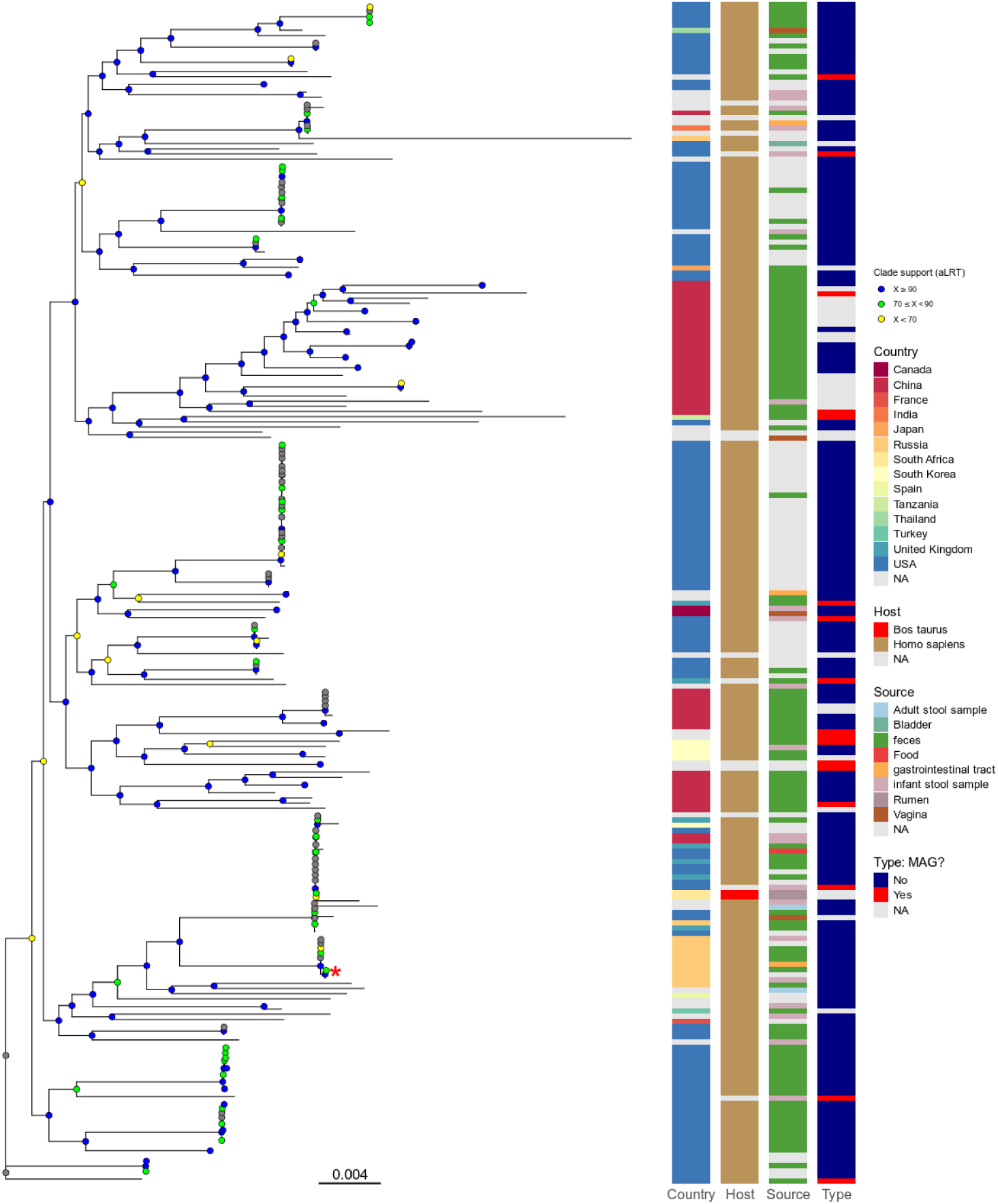
Phylogenomic tree showing the relationships between different *Bifidobacterium bifidum* strains. The tree was created from the alignment of 769 single-copy conserved protein families, using the maximum likelihood method in IQTREE (“-m TEST” mode), with the use of the approximate likelihood ratio test (aLRT) as the branch support test. The outgroup (the proteome of *B. longum* subsp. *longum* JCM 1217) was removed to improve branch resolution. Colored nodes indicate different branch support values ranges. Columns from the right side represent metadata fields reported by the respective NCBI Assembly/Biosample accession. The branch with the red asterisk (“*”) is where the 900791 strain is located.

In order to gain more insights about the genomic patrimony of these species, we formulated a pangenome analysis using the ortholog matrix produced by Orthofinder data, in combination with other tools, to interrogate different aspects of the *B. bifidum* pangenome (Figure 2). The analysis of a total of 414,526 CDS showed that the pangenome of this species is composed of 4,679 orthogroups, 1,278 of them represented by unique genes and other 32 species-specific orthogroups with two or more copies. The core orthogroup set (i.e., the set of ortholog families present in all 229 genomes) contained 1,212 families, 930 of them composed by single-copy orthogroups. Orthofinder data suggests that no orthogroup was unique in the 900791 strain in comparison with the other 228 genomes; interestingly, 900791 shared 23 orthogroups exclusively with the eight genomes from the aforementioned clonal subgroup strains, encoding several ORFans, membrane proteins and a potential lantibiotic biosynthetic cluster (Supplementary Table 2); lantibiotics are well-known compounds that function as antibiotics, causing antimicrobial effects by either forming lethal membrane pores or via inhibition of peptidoglycan biosynthesis (Brötz and Sahl, 2000). The clustering pattern in orthogroup heatmap (Figure 2-A) suggests that the same genomes related to 900791 strain were also related in terms of orthogroup content, increasing the possibility of a clonal subgroup (see above). The pangenome accumulation curve (Figure 2-B) showed a power law equation (*y = ax*^γ^) with a gamma value of 0.21512, indicating that this pangenome is open and the complete accessory of this species is yet to be largely incremented. When the functional annotation for the core/near-core (i.e., gene families with >= 95% prevalence among genomes) gene fraction was compared with the accessory gene fraction (i.e., families with < 95% prevalence), a well-defined conservation profile was obtained (Figure 2-C): functions associated to information transfer (categories J: “Translation, ribosomal structure and biogenesis”) and metabolism (categories F: “Nucleotide transport and metabolism”, C: “Energy production and conversion”, H: “Coenzyme transport and metabolism”, I: “Lipid transport and metabolism”, and E: “Amino acid transport and metabolism”) were found with a high Log2 ratio for the core-vs-accessory fractions, suggesting that those kind of functions have been functionally selected to be part of most *B. bifidum* genomes as possible. Additionally, the strong presence of translation-associated genes in the core fraction, such as the ones involved in the ribosome assembly and structure, may reflect how these genes are recalcitrant to be transferred horizontally, making them less susceptible to be present in the accessory fraction (Kanhere and Vingron, 2009). In counterpart, genes in categories X: “Mobilome: prophages, transposons” and V: “Defense mechanisms”, as well as genes without COG (see the ‘@’ boxplot in Figure 2-C), were found with the more negative Log2 ratios in the core-vs-accessory comparison. The presence of this kind of functions, as well as the enrichment of ORFans in the accessory pangenome, as potential main contributors to the openness of a pangenome, are already well-documented phenomena (e.g., (Rajput et al., 2023)).

**Figure 2.**
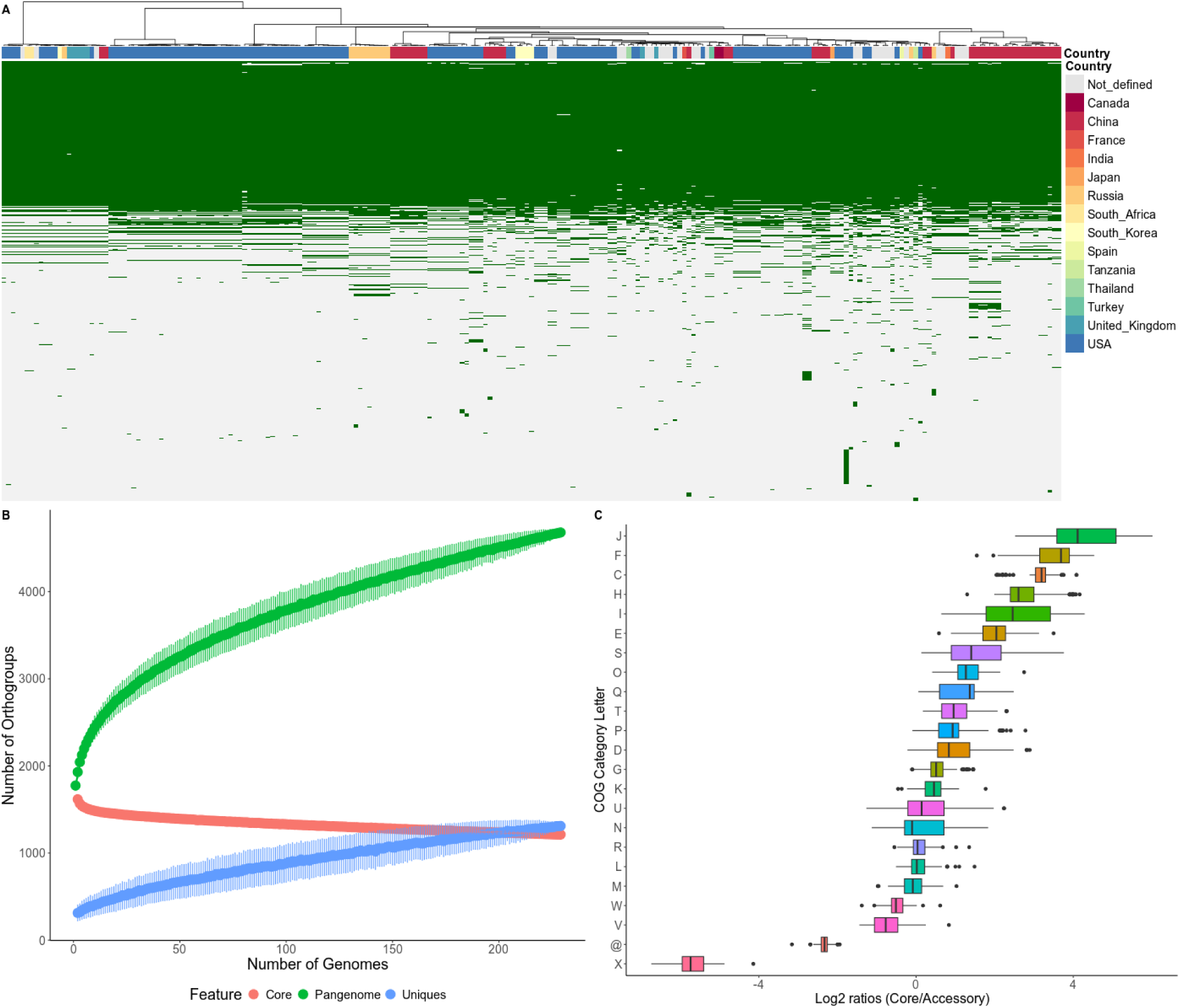
Pangenome analysis of *B. bifidum* utilizing 229 high-quality genomes (including 900791 strain). A: Ortholog conservation heatmap for the genomes according to their geographical origin, clusterized their ortholog profile. B: Pangenome accumulation curve suggests that the pangenome for *B. bifidum* is open (gamma value = 0.21512, according to Power Law equation). C: Analysis for the Log2 ratios of the percentage of core/near-core (>= 95% prevalence) versus accessory genes (< 95% prevalence) genes for each COG category among pangenomes “(@” = no COG category). COG category descriptions are available on the *COG Database website* (https://www.ncbi.nlm.nih.gov/research/cog).

A relevant feature expected to be found in *B. bifidum* strains is the presence of genes involved in the degradation of a variety of glycans and fibers. *B. bifidum* strains harbor a variety of GHs (Figure 3). This feature has been copiously reported in several studies, showing capabilities of different *B. bifidum* to degrade human milk oligosaccharides (HMOs, (Kitaoka, 2012)), mucin glycans (Glover et al., 2022) and some relevant disaccharides such as lactose (Passerat et al., 1995). In the case of HMOs, several previous studies have proposed the use of glycosyl hydrolases from families GH33 (Exo-α-sialidases, EC 3.2.1.18), GH29/GH95 (L-fucosidases), GH2/GH42 (β-galactosidases), GH16 (endo-beta-1,4-galactosidases), GH20/GH84 (β-N-acetilglucosaminidases), GH36/GH129/GH101/GH109 (α-N-acetilgalactosaminidases / endo α-N-acetilgalactosaminidases), GH89 (α-N-acetilglucosaminidasas), and GH36/GH110 (alpha-galactosidases). Interestingly, glycans found in mucin, a protein found in the linings of the gastrointestinal tract, also can be harnessed by different GHs, some of them with common families also required with HMOs (Raba and Luis, 2023). Previous studies showed that *B. bifidum* in general harbors genes encoding GH33, GH29, GH95, GH20, GH2, GH42, GH101, GH129, GH89, and GH84, making this species able to cleaving different glycosic bonds in mucin glycans (Glover et al., 2022). In the case of lactose degradation, bifidobacterial strains can utilize beta-galactosidases from GH2 and GH42 for this purpose (Ambrogi et al., 2019). In order to evaluate these properties from the genomic data of the 900791 strain in comparison with other strains from the species, we searched for CAZymes (using the HMM profile collection from dbCAN-CAZy) and formulated a clustering analysis from their conserved protein patterns (Figure 3). This GH-based clustering also showed the conservation of a subcluster comprising “clonal subgroup” strains (Figure 3-A), suggesting that the CAZyme pattern is also indicative of the common features of this geography-specific group. The main GHs collaborating with data dispersion causing this clusterization were associated (according to their entries in CAZYy) to the following activities: β-galactosidase (GH42), xylan exo/endo-β-1,4-xylosidase (GH43_2, GH43_10, GH43_35), β-galactofuranosidase (GH43_30, GH43_32), β-glucosidase (GH43_34), different types of α-L-arabinofuranosidase (GH43_5, GH43_9), b-1,3-lacto-N-biosidase (GH136) and others (Figure 3-B). In the particular case of the 900791 strain genome, the following GHs were detected: GH1, GH101, GH109, GH110, GH112, GH121, GH129, GH13, GH136, GH16_8, GH164, GH167, GH177, GH179, GH184, GH188, GH2, GH20, GH23, GH25, GH29, GH3, GH30_5, GH32, GH33, GH35, GH36, GH42, GH43, GH5, GH51, GH77, GH84, GH89, GH93 and GH95. From these families, groups such as GH2, GH20, GH33 and GH84 have been previously identified in mucin (Glover et al., 2022) and HMO degradation (Lordan et al., 2024). The detection of these GHs in the genome of 900791 suggests that this strain is also to degrade these kinds of compounds. Likewise, GHs from families GH2 and GH42 with a potential role in lactose degradation were found in the genome of 900791. The detection of these candidates has agreement with the previous observation in which this strain helped to the improvement of lactose tolerance in human subjects (Aguilera et al., 2021).

**Figure 3.**
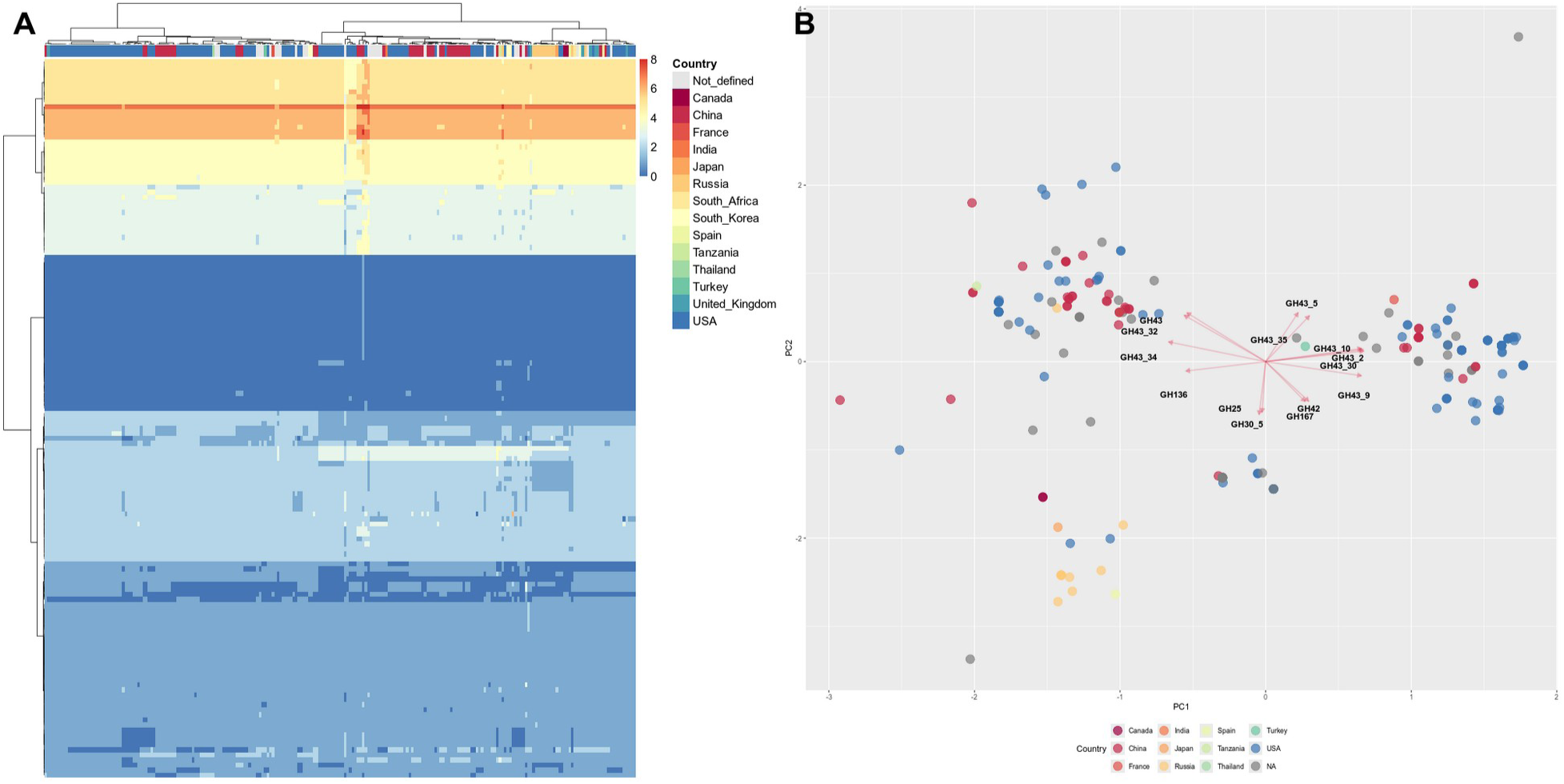
Comparative analysis of Glycosyl hydrolases (GHs) profile analysis of *B. bifidum* utilizing 229 high-quality genomes (including 900791 strain), characterized by country of origin. HMM profiles from the dbCAN version 15 were used against the ORFomes of 229 *B. bifidum* strains and clusterized (by euclidean distances) in a heatmap (A) and in a PCA dotplot showing the most important loadings influencing the dispersion of the genomes by their GH content (B).

### 3.3. Predicted probiotic features of the 900791 strain and its clonal subgroup

As seen previously, the 900791 strain is related with other *B. bifidum* strains forming a potentially clonal group. Since this strain is an interesting probiotic candiate, it is necessary to track such potential features in its genome to predict its success and utility as a probiotic agent. Genomic information available for different probiotics are a valuable source for the search and identification of genotypic traits that ensure efficacy, safety, and stability. Strain-specific genetic variability requires a precise understanding of conserved and accessory genes to optimize selection criteria. According to previously studied models, some important markers for probiotic properties include genes involved in adhesion to intestinal mucosa, polysaccharide synthesis, and substrate utilization, which are essential for gut colonization and persistence (Rossi et al., 2022). Stress factors involved in low pH and bile salt resistance are also important features to search in probiotics (Ruiz et al., 2013; Collado and Sanz, 2007). Additionally, other desirable traits are the verified absence of antimicrobial resistance markers and virulence factors (Li et al., 2024) and the ability to exhibit a proper oxidative stress response (Feng and Wang, 2020).

In order to establish predicted features of the 900791 strain and its related strains, we will separate this section into different features:

#### 3.3.1. Adhesion to mucin and epithelial cells

Different studies have found proteins involved in the binding of *Bifidobacterium* to mucin and the intestinal epithelium, a useful feature for probiotics to exert their beneficial health effects, such as strengthening the intestinal barrier, preventing pathogen colonization, and influence on the immune system of the host (Monteagudo-Mera et al., 2019). In *Bifidobacterium*, this feature has been studied: in *B. bifidum* PRL2010 strain, sets of sortase-dependent pili (FimA-FimB proteins, encoded by multiple gene copies) were involved in adhesion to epithelial cells (Milani et al., 2017). Another study, performed on *B. bifidum* BBMN68, showed a novel protein, called FimM, involved in the adhesion of *B. longum* to mucin, fibronectin, and fibrinogen (Xiong et al., 2020). Both the FimA/FimB and FimM proteins are members of the COG4932 family, involved in bacterial binding to different matrices and cells. Interestingly, in the 900791 genome and its clonal group, at least three orthogroups belonged to COG4932, suggesting that this strain and its related genomes contains cell-binding abilities by the use of FimA/FimB and FimM proteins (Figure 4A).

**Figure 4.**
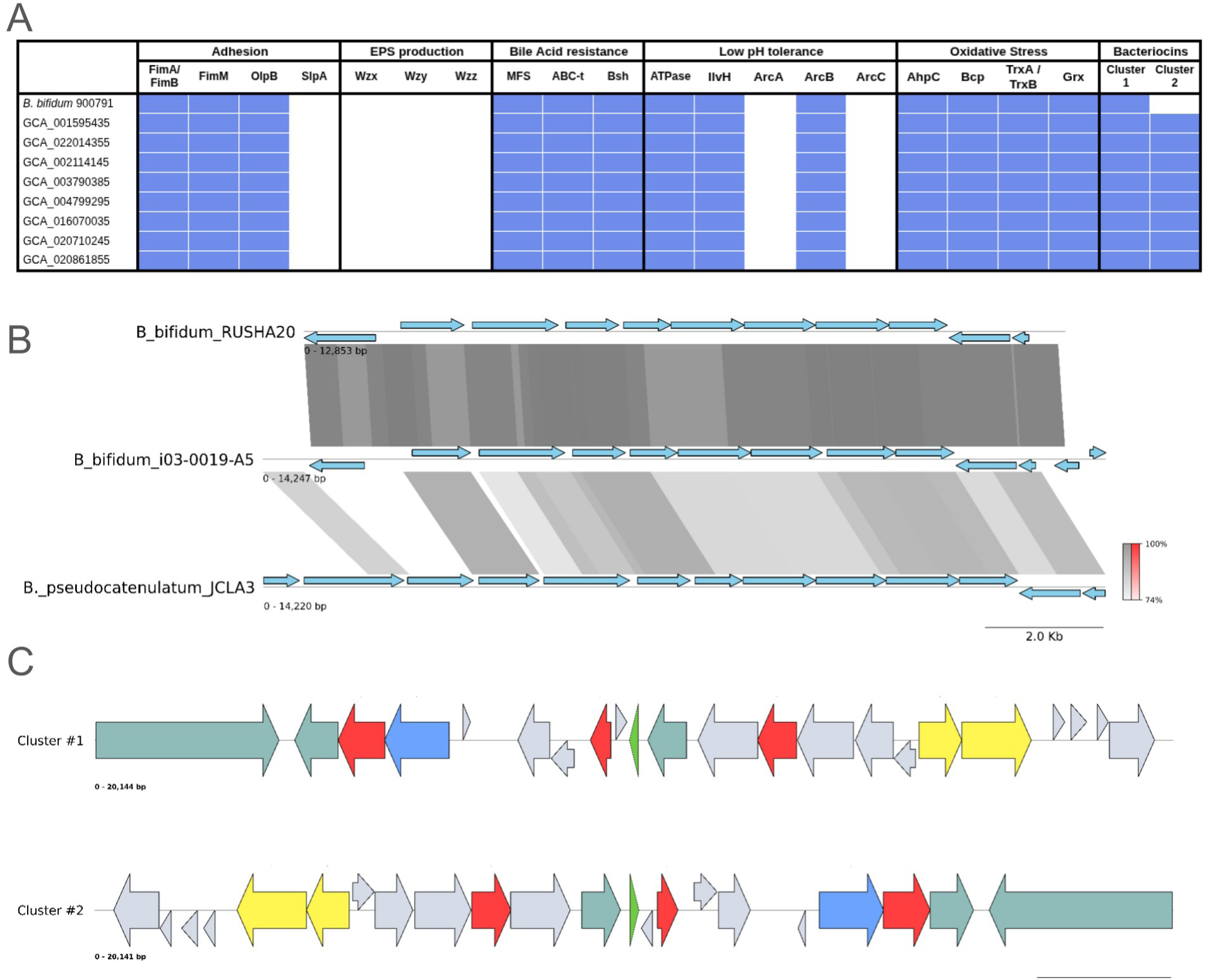
Conservation of genes involved in A: table of genetic features searched in the characterization of the 900791 strain and the other members of the clonal group. B: Conservation of the closest gene cluster potentially involved in EPS biosynthesis. C: features of the found clusters/AOIs involved in bacteriocin biosynthesis, showing relevant proposed roles in different colors; Green: bacteriocin-encoding gene; red: genes encoding components of ABC-type transporters; yellow: potential regulators, dark green: potential multidrug exporting systems; blue: genes involved in lantibiotic biosynthesis; grey: other functions.

Another potential bacterial structure involved in adhesion is the proteinaceous surface layer (S-layer). A study in different *Lactobacillus* strains (Frece et al., 2005; Prado Acosta et al., 2016) showed that S-layer is responsible for adhesiveness of epithelial cells, bacterial protection and even interaction with receptors from host cells. A previous study has suggested the presence of S-layer proteins in bifidobacteria (Li et al., 2018) but the functionality of those proteins in *Bifidobacterium* are scarcely studied yet. At least, our *in-silico* prediction for the 900791 strain and its clonal group suggest the presence of a gene encoding a member of the COG1361 family; this family also contained a protein called OlpB, a surface protein associated with the cellulosome in *Clostridium thermocellum* and potentially involved in direct interaction with cell surface (Fujino et al., 1993). Although this protein is not similar to the classic SlpA S-layer proteins found in *Lactobacillus (Boot et al., 1993)*, we hypothesize that it may have a similar role (Figure 4A).

#### 3.3.2. Exopolysaccharide Production

Another relevant feature associated with microbe-host interactions in *Bifidobacterium* is the ability to produce exopolysaccharides (EPSs), commonly produced polymers composed by repeated mono- or oligosaccharide units connected by glycosidic links. There are studies supporting the idea that surface-associated EPSs support *Bifidobacterium* gut colonization (Sadeghi et al., 2024); moreover, comparative genomics studies showed that several *Bifidobacterium* strains contained at least one copy of a cluster (the eps cluster) potentially involved in EPS production, even if in several *B. bifidum* strains, this potential gene cluster was absent (Ferrario et al., 2016). In general, an *eps* cluster contains genes for A) the priming enzyme that catalyzes the initial step of EPS-unit synthesis via the transferring of a sugar phosphate to a lipophilic carrier molecule anchored to the cell membrane; b) different glycosyltransferases and enzymes involved in the metabolism of nucleotide sugars and, c) an export-polymerization system, generally containing flippases (Wzx), polymerases (Wzy) and a chain-length determination protein (Wzz, Hidalgo-Cantabrana et al., 2014). From these compounds the priming enzyme and the export/polymerization system are the most distinctive. Our analysis of the 900791 strain genome showed only one potential cluster (surrounded by transposases) involved in the production of some EPS precursors, glycosyltransferases and a putative ABC transport system (Figure 4B); this gene cluster shares high nucleotide identity (> 99%) with a similar cluster found in the the assembly of *B. bifidum* i03-0019-A5 (accession AP031420), and moderately high identity (74% approximately) with a cluster from *Bifidobacterium pseudocatenulatum* JCLA3 (accession CP090598). However, it was concerning that our strategy failed to find potential candidates for a proper export-polymerization system, including a flippase, a polymerase and a chain-length determination protein; even using the previously reported sequences from other bifidobacteria, these components could not be found in the genome of 900791 strain and its clonal group. This information may suggest that this strain may not produce EPS, and this affirmation remains to be supported by experimental studies.

#### 3.3.3. Bacteriocin production

Another important feature to be observed in probiotics is the ability to produce bacteriocins, peptides that are active against other bacteria without causing effect on the owner (Anjana and Tiwari, 2022); additionally, some bacteriocins are not only associated to antimicrobial activity, but they can also function as signalling peptides involved in quorum sensing or interspecies communication (Dobson et al., 2012). Therefore, bacteriocins may offer a competitive advantage to probiotics in their passage or colonization in the gut. By using BAGEL, we searched for bacteriocin-associated loci in the genomes of the 900791 strain and its clonal group. This search could find two potential gene clusters (also called as “areas of interest” or AOIs) encoding bactericins in the members of the clonal group of the 900791 strain (Figure 4C); curiously, only one of the clusters was found in the 900791 strain itself. Cluster/AOI #1 (detected in the 900791 strain and practically identical in the other strains) contained a potential bacteriocin related to the *Geobacillin_I_like* family, a member of the Lanthipeptide A subclass of bacteriocins. Representatives of this family are the geobacillins found in *Geobacillus* species, with experimentally validated antimicrobial activity on different bacterial species (Garg et al., 2012). Cluster/AOI #2 (not detected in the 900791 strain, but found in the eight other members of the clonal group), contained a putative bacteriocin from the *Propeptin_2* family of the *Lasso peptide* subclass. The representative of this aforementioned family is a peptide found in *Microbispora* sp. SNA-115, with inhibitory effect on prolyl endopeptidases and validated antimicrobial activity against *Mycobacterium phlei* (Kimura et al., 2007). In both clusters/AOIs, some genes encoding components of ABC transporter systems or genes involved in lantibiotic biosynthesis (containing domains PF05147 or PF00733). Thus, these predictions suggest that 900791 may contain functional bacteriocins with potential impact on different bacteria.

#### 3.3.4. Acid (low pH) resistance

One desirable trait in potential probiotic microorganisms is the ability to tolerate low pH, since it is required for survival in their passage in the acidic gastric environment (Huys et al., 2013). Previous studies have shown that bifidobacteria contain several physiological adaptations to low pH resistance, and those are more effective under pre-acclimation (Jin et al., 2015, 2012). Despite several phenotypic traits are associated to acid stress in *Bifidobacterium*, some key genetic markers include the F₀F₁-ATPase system (*atp* genes) for proton extrusion and branched amino acid biosynthesis (Sánchez et al., 2007); in some organisms such as *Lactobacillus sakei*, arginine deimination is part of the low pH responses, by the use of a three-step pathway (*ArcA*, arginine deiminase; *ArcB*, ornithine carbamoyltransferase; *ArcC*, carbamate kinase) resulting in the conversion of arginine into ornithine, ammonia (compound that works as proton acceptor), and carbon dioxide (Zúñiga et al., 1998; Rimaux et al., 2012). The analysis of the 900791 strain and its related genomes showed that all nine strains contained genes for a potentially functional F₀F₁-ATPase system, suggesting that these strains may also utilize this strategy. A phenotypic study already showed that ICS-310, one of the strains strongly related to 900791 in its “clonal group”, was already evaluated in their gastric-like acid tolerance, showing a promising resistance phenotype (Perunova et al., 2025). In these 9 genomes, a copy of the IlvE protein (trEMBL accession I3WHR6) encoding a branched-chain-amino-acid transaminase (EC 2.6.1.42), an enzyme involved in branched amino acid biosynthesis; this enzyme is proposed in studies in *Streptococcus mutans* as part of a low-pH metabolic response (Len et al., 2004) and it was also its upregulation was observed in *Bifidobacterium longum* under low-pH stress (Sánchez et al., 2007). In the case of the arginine deiminase pathway, in the nine genomes only one copy of the Ornithine carbamoyltransferase (*ArcB*) was found (using the O53089 trEMBL accession). No related sequences to the arginine or agmatine deiminase enzymes (directly involved in ammonia release) were found, suggesting that this pathway may not be utilized by these strains of *B. bifidum* (Figure 4A).

#### 3.3.5. Bile Salt Resistance

Bile salts are one of the common stress-inducing factors that microorganisms must face in the gastrointestinal tract in their colonization or passage process. The ability to resist those salts is one desirable feature to address in potential probiotic strains (Huys et al., 2013) since this feature may ensure the ability of the probiotic to pass through the gastrointestinal tract. In the case of members of the *Bifidobacterium* genus, several genes have been identified as determinants of bile salt resistance. For example, in *Bifidobacterium breve* UCC2003, the gene Bbr_0838 (Uniprot accession) was found to encode a membrane protein related to MFS transporters, highly induced by bile salts; furthermore, disruption of this gene resulted in a decreased survival in the presence of bile salts, suggesting its role as a resistance protein (Ruiz et al., 2012a). Previous to this finding, a potentially homologous gene to Bbr_0838, found in *Bifidobacterium longum* NCC2705, known as BL0920 exhibited similar structural and phenotypic features (Gueimonde et al., 2009). Another study, also performed in *Bifidobacterium breve* UCC2003, suggested that other transporters from the ABC family (identified as Bbr_0406-0407, Bbr_1804-1805, and Bbr_1826-1827), can also mediate bile salt resistance via extrusion (Ruiz et al., 2012b). In *Bifidobacterium*, Bile salt hydrolases (BSH) are also described as enzymes that break down conjugated bile salts and contribute to tolerance; the case of the *bsh* gene, early found in *Bifidobacterium longum* SBT2928 (Tanaka et al., 2000) is a protein capable to hydrolyze a variety of human and animal bile salts. This gene was also conserved in other bifidobacteria (Schöpping et al., 2022a). The search of these factors in the 900791 and its clonal group found also potential representatives for at least, the MFS transporter Bbr_0838, genes for the ABC subunits (Bbr_0406-0407, Bbr_1804-1805) and for the *bsh* gene, suggesting that this response may utilize both enzymatic and extrusive strategies to resist bile salts in this clonal group (Figure 4A).

#### 3.3.6. Oxidative Stress

Another desired feature to search in probiotic strains is the presence of proteins involved in oxidative stress response (Schöpping et al., 2022b; Imlay, 2003). Since probiotics need the ability to survive during industrial production and storage, as well as their transit through the gastrointestinal tract, the response to sudden changes in reactive oxygen species (ROS) is key in their successful maintenance, passage and survival (Blazheva et al., 2022). Therefore, this type of response is important to aim for in probiotics analysis. In the response to oxidative stress, essential genes include those encoding the thioredoxin-thioredoxin reductase system (TrxA-TrxB), NADH oxidases, superoxide dismutase, catalases, rubrerythrins, and different types of peroxidases or peroxiredoxins (Lu and Holmgren, 2014; Imlay, 2003). Additionally, enzymes involved in the use of different thiol-active compounds (e.g., glutathione) such as glutaredoxins are also markers of oxidative stress tolerance (Averina et al., 2023). The search of proteins associated to oxidative stress response in the 900791 strain and its clonal group found representatives for different well-established proteins such as the Alkyl hydroperoxide reductase (AhpC peroxiredoxin, COG0450), glutaredoxins (Grx, COG0695), Peroxiredoxins (Bcp, COG1225), thioredoxin - thioredoxin reductases (COG0526: TrxA; COG0492: TrxB) (Figure 4A). Several of those proteins are associated in the ROS scavenging and redox rescue of different substrates. This prediction suggests that the 900791 strain may have the ability to tolerate certain levels of ROS exposition.

## 4. Discussion

In this study, we analyzed the complete genome of *Bifidobacterium bifidum* strain 900791 and we compared its ORFeome with the content of 228 other *B. bifidum* genomes; from this study. several important insights emerge regarding its potential as a probiotic candidate and its distinctive genomic characteristics. This discussion evaluates the strain’s properties within the broader context of *B. bifidum* biology and probiotic functionality. The genomic characteristics of strain 900791 are consistent with typical *B. bifidum;* for example, its genome size, number of ORFs and %G+C fall within the expected range for *B. bifidum*, as previous studies have reported: genome sizes: 2.03–2.55 Mb (average = 2.17 ± 0.09 Mb; average of 1837 ± 143 ORFs per genome; G+C content ranging between 62.3% and 62.8% (Lu et al., 2021). The functional annotation reveals a genome adapted for gut colonization and carbohydrate metabolism, hallmarks of successful gut commensals, seen commonly in other members of the species (Glover et al., 2022), (Raba and Luis, 2023). The presence of a rich variety glycosyl hydrolases (GHs) from families associated with degradation of HMOs and mucin glycans, including GH2, GH20, GH33, and GH84, underscores the capacity of this *B. bifidum* strain for utilizing host-derived glycans, in correspondence with the fact that this strain was obtained from human feces. Additionally, the detection of GH2 and GH42 families specifically associated with lactose degradation provides genomic support for the previously observed clinical improvement in lactose tolerance in human subjects consuming this strain (Aguilera et al., 2021). The subcellular localization predictions indicating that 28.2% of proteins are located outside the cytoplasm suggest an active secretome and membrane-associated machinery that may reflect the great importance of host-microbe interactions for the physiology of this organism, covering secretable CAZymes, active proteins and other components influencing the interaction with the host-associated environment (Vidal-Veuthey et al., 2022). Thus, this extensive repertoire of exposed proteins is consistent with the lifestyle of gut-adapted bifidobacteria, which must interface effectively with the host intestinal environment.

This study, as far as we know, also presented the biggest *B. bifidum* pangenome analysis to date; the open pangenome structure may indicate the need of genome plasticity to adapt to different contexts in the human gut: the requirement for an ongoing gene content change is mostly impulsed by changing niches, diverse ecological interactions and large effective population sizes (Brockhurst et al., 2019). Due to these reasons, the open pangenomes are a common feature among gut bacterial species (Shoer et al., 2024). In this context, the functional enrichment patterns observed core fraction is completely expectable: genes involved in translation, amino acid metabolism, and energy production, reflect the essential cellular processes required for survival. The overrepresentation of accessory genes in mobilome and defense functions indicates that these represent major sources of genomic variation and adaptation within the species; this observation was made in other bacterial organisms (e.g., (Liao et al., 2023)).

The phylogenomic analysis, in combination with cgMSLT analysis, showed that 900791 belongs to a distinct clonal subgroup comprising nine closely related strains, all isolated from human stool samples (mostly from children). This clonal relationship, supported by >99% allelic similarity in cgMLST analysis, suggests recent common ancestry and shared adaptive strategies; also, the geographical clustering of this clonal group was relevant (all strains came from the same country, see Figure 1); these patterns suggest that possible population structure within *B. bifidum* may reflect local adaptation or founder effects. Moreover, the identification of 23 unique orthogroups exclusive to this clonal subgroup (including a set of genes putatively involved in lantibiotic biosynthesis), reinforce the importance of these strain-specific adaptations.

Very importantly, this study provided the *in-silico* characterization of the potential use of the 900791 strain as a probiotic. The safety profile of 900791 represents a pivotal foundation for the justification of its probiotic potential. The comprehensive antibiotic resistance analysis using three independent methods (NDARO, ResFinder, and RGI-CARD) revealed only intrinsic resistance to mupirocin and rifampicin, mediated by naturally occurring mutations in *ileS* and *rpoB* genes respectively. These resistance markers are well-documented common mutations in bifidobacteria and those strains have been used as probiotics regardless of the presence of those resistance markers (Grmanová et al., 2010; Serafini et al., 2011). Since determinants for acquired or transferrable antimicrobial resistance genes, plasmids, and virulence factors were not found, it is possible to suggest that this strain has a safe antimicrobial resistance profile for a probiotic.

The genomic analysis also revealed multiple stress tolerance systems that are essential or desirable for probiotic functionality: genes involved in the tolerance to low pH experienced during gastric passage (e.g., by the presence of the F₀F₁-ATPase systems or a branched-chain amino acid transaminase); genes involved in the extrusion or conversion of bile salts to improve the tolerance to those compounds in the passage by the small Intestine (by encoding Bbr_0838-like MFS transporters, ABC transporter subunits, and the Bsh hydrolase); or the potential ability to tolerate fluctuant exposition to ROS via a proper oxidative stress response (via enzymes such as thioredoxin-thioredoxin reductase systems, alkyl hydroperoxide reductase, peroxiredoxins, glutaredoxins). All those stress responses predicted in the genome of the 900791 strain and conserved in the rest of its clonal group may suggest that this strain may be well equipped to the challenges that its use as a probiotic requires.

The genomic repertoire of 900791 also included several factors associated with beneficial host-microbe interactions: the identification of genes encoding FimA/FimB and FimM, as well as a putative S-layer protein, indicates the potential of this strain to interact with mucin or epithelial cells, in order to improve colonization efficiency, another desired feature for probiotic bacteria. Additionally, bacteriocin production capability of 900791 (and also for the other strains of their clonal group) represents a significant competitive advantage in the complex gut ecosystem. This potential antimicrobial capacity may contribute to the ability of 900791 to suppress pathogenic bacteria, a desirable probiotic trait (Yu et al., 2023).

Finally, considering that whereas genomic analysis provides strong evidence for the probiotic potential of 900791, experimental validations of the predicted stress tolerance mechanisms under physiologically relevant conditions, or for the other features, are key aspects to work in the future. In-vitro (and further in-vivo) studies evaluating colonization dynamics, persistence, and interaction with the existing gut microbiota would provide crucial insights into the key features of the 900791 phenotype as a candidate probiotic. Moreover, clinical trials evaluating specific health benefits beyond lactose tolerance, such as immune modulation or pathogen exclusion, also are important aspects to define in order to gain a better insight of the therapeutic potential of this strain. Experimental, comparative studies with other members of the clonal subgroup could also reveal whether the shared genomic features translate to similar probiotic properties, potentially identifying a group of related strains with complementary or enhanced therapeutic applications.

## 5. Conflict of Interest

A.G is the Chief Executive Officer of *Bifidice SpA*. Authors D.A, B.V-V and J.P.C served as advisors in *Bifidice* SpA and were contacted by A.G to conduct the analysis of this study. K.M holds a researcher position as an employee of *Bifidice* SpA.

## 6. Author Contributions

J.P.C and A.G conceived and directed the study; J.P.C, B.V-V and K.M analyzed the data; all the authors collaboratively elaborated, edited, and corrected the text and figures in the manuscript. All the authors read the paper and approved the content.

## 7. Funding

B.V-V is supported by ANID Doctorado Nacional/2021-21211564. This research was partially supported by the computing infrastructure of the *Centro de Genómica y Bioinformática*, *Universidad Mayor*.

## Supplementary Data

**Supplementary Table 1.** List of the 228 *B. bifidum* genomes retrieved from NCBI and selected for this study, including information from GTDB and CheckM2 results, and metadata retrieval.

**Supplementary Table 2.** List of the functional annotation of the set of 23 orthogroups exclusively conserved in the clonal group related to the 900791 strain.

**Supplementary Figure 1.** Circular representation of the complete genome of the 900791 strain including genomic features and predicted functions according to COGs.

**Supplementary Figure 2.** Main annotation stats for the 900791 strain ORFeome. A: Piechart showing the predicted subcellular localization according to PSORTb 3.0 prediction (Gram-positive mode). B: Piechart showing the distribution of the COG metacategories according to EGGNOG mapper predictions. C: barplot showing the represented COG categories in the ORFeome.

