## Supplementary figures and images for "Genomic analysis of the probiotic candidate *Bifidobacterium bifidum* strain 900791 in the context of the *B. bifidum* pangenome"

### Supplementary Figure 1

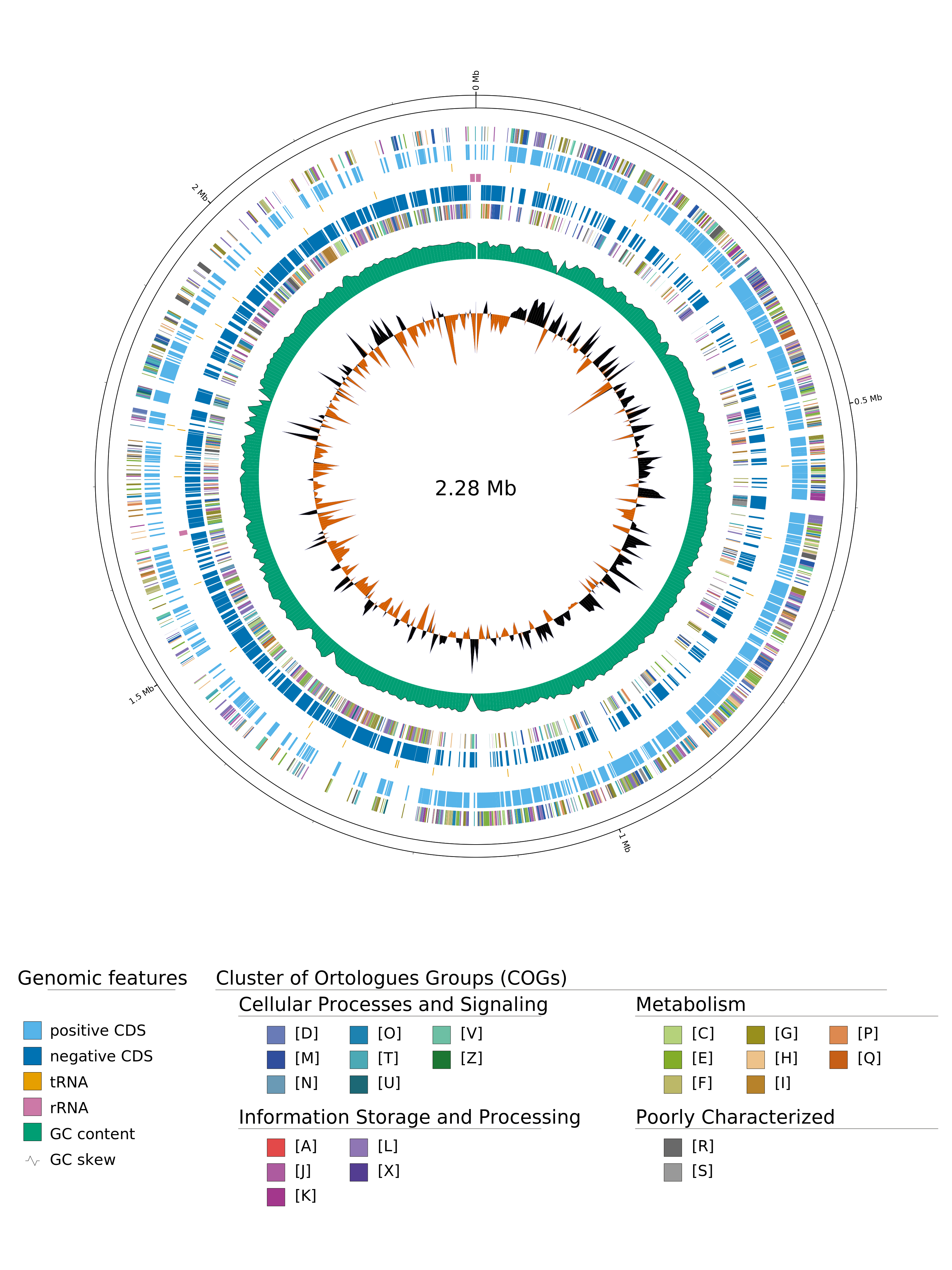

### Supplementary Figure 2

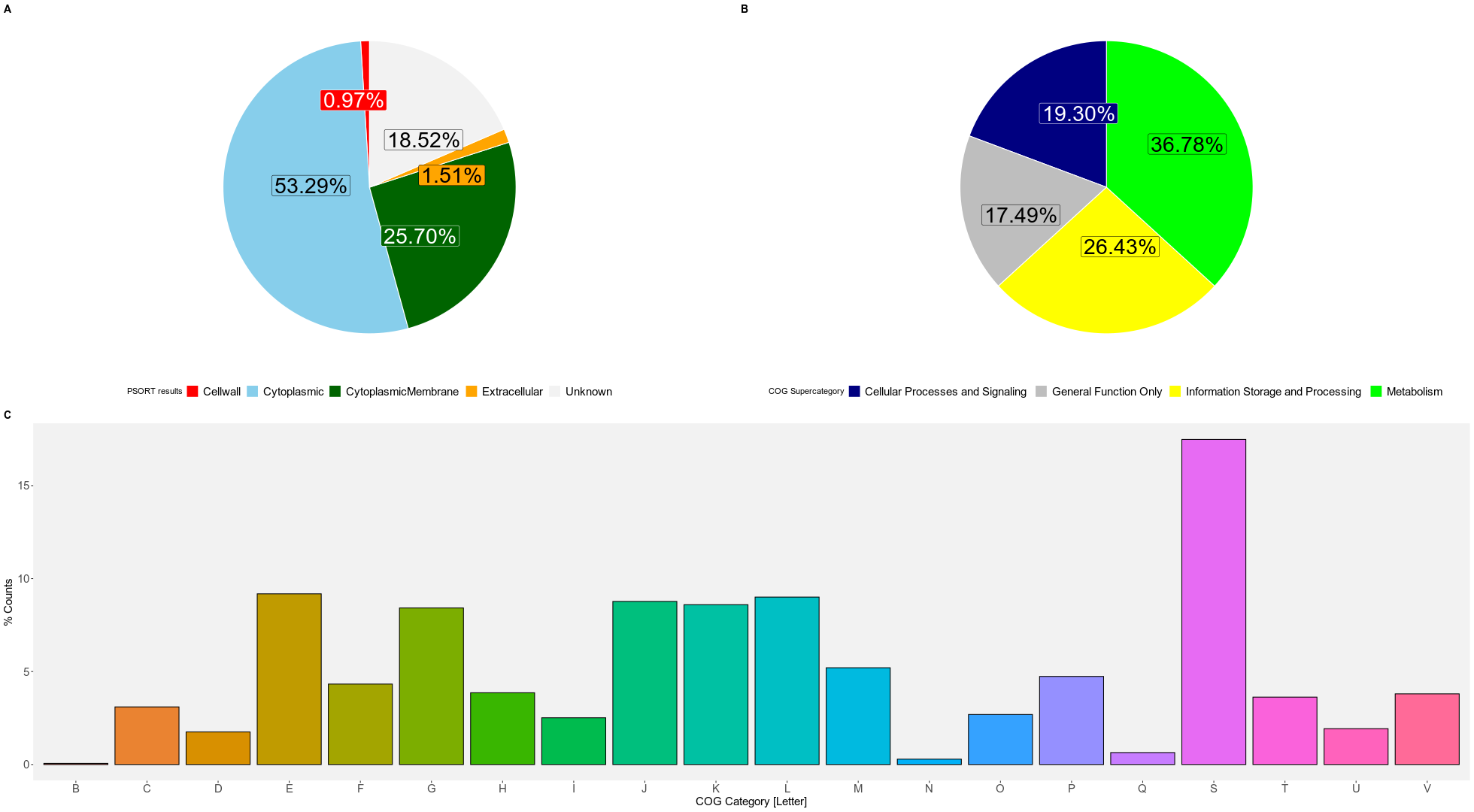
